# Fractured tonotopy of functional neural clusters in mouse auditory cortex

**DOI:** 10.1101/059493

**Authors:** Isabel Delgado Ruz, Simon R. Schultz

## Abstract

The degree of order versus randomness in mammalian cortical circuits has been the subject of much discussion. Previous reports showed that at a large scale there is smooth tonotopy in mouse auditory cortex, while at the single neuron level the representation is the traditional “salt and pepper” configuration attributed to rodent cortex. Here we show that at the micro columnar scale we find a large variety of response profiles, but neurons tend to share similar preference in terms of frequency, bandwidth and latency. However, this smooth representation was fractured and large differences were possible between neighbouring neurons. Despite the tendency of most groups of neurons to operate redundantly, high information gains were achieved between cells that had a high signal correlation. Connectivity between neurons was highly non-random, in agreement with a previous in-vitro report from layer five. Our results suggest the existence of functional clusters, connecting neighbouring mini-columns. This supports the idea of a “salt and pepper” configuration at the level of functional clusters of neurons rather than single units.

## 1. Introduction

The topographic organisation of the mammalian cortex has long been a topic of research, with the presence of orderly maps (or the lack thereof) in some cortical areas being a key difference between species. Examples of such topographic organisation include tonotopy in the auditory cortex, retinotopy in the visual cortex, and orientation columns in the visual cortex of some mammals. Mouse cortex has often been given as a typical example of a random arrangement with its “salt and pepper” configuration. The organization of mouse auditory cortex has however recently been under debate, with two photon calcium imaging studies showing heterogenous organisation in terms of preferred frequency when considering spiking activity from neighbouring neurons from upper layers, while showing tonotopy when considering the aggregated activity of groups of neurons (Rothschild et al., 2010; Bandyopadhyay et al., 2010). Recent electrophysiological studies have shown that tonotopy largely depends on the type of signal considered for the characterisation, quality of tuning included, area of auditory cortex, anaesthetic state and layer with upstream layers having a degraded tonotopy compared to thalamo-cortical recipient layer (Hackett et al., 2011; Guo et al., 2012).

We can expect that any property exhibited by a population of neurons must have an underlying biophysical correlate. It has been proposed that local connectivity plays an important role in shaping the arrangement seen in cortices lacking in orderly arrangement of columns as is *Preprint submitted to Elsevier* the case with mouse cortex, while the existence of structure such as pinwheels and the columnar arrangement around them is mainly due to long range suppressive connections (Kaschube et al., 2010). Experimental evidence from in-vitro studies has shown mini columnar ensembles (Maruoka et al., 2011; Krieger et al., 2007) not yet been characterised in vivo, which in principle could aggregate to form functional units as suggested by (Krieger et al., 2007) and (Ohki et al., 2005) and could constitute the basis of columnar processing by sharing common input and being synaptically connected.

In this study, using high density columnar micro electrode arrays, we measured functional properties of local columnar populations from layer V. Reconciling the known random configuration exhibited by mouse cortex and the columnar processing expected to take place, we found that groups of neighbouring neurons tend to share similar properties while still having a rich variety of response profiles, coherent with the existence of overlapping fine-scale networks (Yoshimura and Callaway, 2005; Yoshimura et al., 2005). We studied monosynaptic interactions within these local networks, finding highly non-random connectivity and their spatial coverage supports the idea of inter column functional aggregation.

## 2. Methods

### 2.1. *Surgery and Stimuli*

We performed experiments on 5-7 week old mice (n= 12 CBA/Ca mice) under terminal anaesthesia, in strict ac-cordance with the 1986 Act (Scientific Procedures) under license granted by the UK Home Office. Mice were deeply anaesthetised with a mixture of fentanyl (0.05 mg/Kg), midazolam (5 mg/Kg) and medetomidine (0.5 mg/Kg). The animal’s temperature was maintained at 37°C using a feedback controlled blanket.

A small craniotomy was performed over auditory cortex area A1, and recordings with high density electrode arrays were made using a Poly3-25s probe (Neuronexus Technologies) as shown in Fig. 1A. The probe was introduced parallel to the midline and lowered to a single location targeting layer 5, collecting both spontaneous and evoked activity.

We used iso-intensity pure tone stimuli (70 dB) generated using an RZ6 real-time processor and presented via electrostatic speaker (ES-1; Tucker-Davis Technologies) placed 5 cm behind the contralateral ear. Pure tones (3-48 kHz, 1/3 octave steps, 100 ms duration), gated with ramped cosine windows (3 ms to 90% of maximum), were presented at a frequency of 0.5 Hz.

### 2.2. *Single unit characterisation*

Single neurons were isolated and localised as described previously (Delgado-Ruz and Schultz, 2014) from raw data collected across the 32 channels as seen in Fig. 1B. Briefly, signals were spike sorted using Caton/MaskedKlustakwik (Kadir et al., 2013), and then each isolated neuron was fitted to a line source model to estimate its location. Examples of sorted neurons can be seen in Fig. 1C and a raster plot of all recorded neurons in Fig. 1D. Each neuron was also classified as narrow spiking or broad spiking (putative inhibitory and excitatory respectively) following the procedure described in (Peyrache et al., 2012), which is based on the temporal waveform of the action potential.

The tuning of each unit was calculated for the 40 ms after stimulus onset. This time window was selected as the responses measured when using larger time windows contain more noise (Moshitch et al., 2006). In addition, upstream neurons are likely to use the first few tens of milliseconds from the neurons’s responses (see Cohen and Kohn (2011) for discussion).

The neuronal response to stimuli was characterised using three metrics: preferred frequency, bandwidth and peak latency from onset. The bandwidth measures the selectivity exhibited by each neuron, while the latency reflects differences on upstream path and distance from soma to synaptic inputs (Carrasco and Lomber, 2009; Bizley et al., 2005; Sakata and Harris, 2009; Katona et al., 2012). Preferred frequency was defined as the frequency that elicited the largest evoked response. Bandwidth was defined as the 3 dB bandwidth around the preferred frequency, and peak latency from onset was defined as the time elapsed from onset until the peak firing rate for the preferred frequency.

Differences in response properties were shown qualitatively using Voronoi diagrams. To construct them the xy cartesian plane is divided into regions (one per neuron) whose boundaries are defined based on neurons’; locations calculating boundaries equidistant to neighbouring neurons. By assigning to each region the property value measured for the neuron associated to it, the Voronoi diagrams allow us to graphically show how the xy plane spans different property values.

The temporal response profile was characterised, identifying neurons that showed ON (transient or sustained response) and OFF responses. This simple separation allowed us to see the range of responses. We used the method described in detail in (Willott et al., 1988), which is based on the computation of an adaptation ratio *A*_*r*_ that quantifies variations in firing rate on the time slots after stimulus onset. Neurons with *A*_*r*_ < 0.2 were considered as having a transient or phasic response, while 0.2 < *A*_*r*_ < 2 were considered sustained. Neurons having a firing rate above 25% above spontaneous level after stimulus offset were considered to have an OFF response.

### 2.3. *Characterisation of interactions*

Each recorded pair was analysed to detect putative monosynaptic connections (Fujisawa et al., 2008), with crosscorrelograms generated as described by Kohn and Smith (2005). Pairwise signal correlation allows the measurement of similarity between the response to stimuli; we calculated it, according to standard usage, as the Pearson correlation coefficient between mean responses to stimuli:

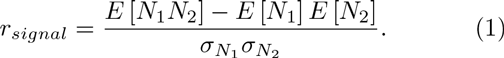

Here *E* is the expected value and *σ* the standard deviation of the mean spike count *N*_*i*_. Noise correlation or spike count correlation measures the tendency of neurons to covary their firing rate, and is calculated as the Pearson correlation coefficient between variations around the mean responses to stimuli (Bair et al., 2001; Kohn and Smith, 2005):

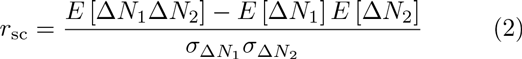
where Δ*N*_*i*_ corresponds to the difference between the mean response of cell **i** to a particular stimulus and its response on a particular trial. Signal correlation may reflect shared inputs driven by the stimuli, whereas noise correlation may reflect slower network oscillations and synchronous activity not linked to stimuli (Cohen and Kohn, 2011). The All-Way Shuffle Predictor and Jitter Corrector were used to correct the crosscorrelograms (Kohn and Smith, 2005; Smith and Kohn, 2008). The shuffle predictor allows the removal of correlations locked to the stimulus, while the jitter corrector removes correlations arising from slow oscillations and stimulus-locked correlations. The All-Way Shuffle Predictor (Kohn and Smith, 2005) corresponds to the CCG calculated after shuffling the id of the different trials, normalised by the mean firing rates, λ_*i*_:

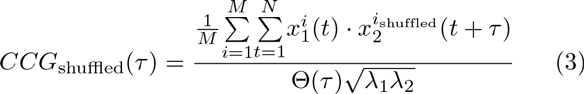
where M is the number of trials, N is the number of bins ⊝(τ) per trial and is a triangular function which corrects for the amount of overlap between spike trains. The Jitter Corrector, see (Smith and Kohn, 2008) for details, corresponds to the CCG calculated over surrogate spike trains. The method for generating the surrogates can be summarised as follows:

- A window for jittering *T*_*j*_ ms is defined.
- Each spike train for each trial is divided in time windows of length *T*_*j*_. The timestamps of all spikes across trials within each time window are added to a pool of spikes.
- On each trial, for each time window, the spikes are replaced for a spike randomly chosen from the corresponding pool of spikes.

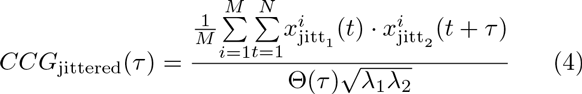

The timescales of correlations, *r*_*ccg*_ (Kohn and Smith, 2005; Bair et al., 2001), were characterised to study di.erence in noise correlations found between this and previous studies, as this metric allows to study the correlations over varying time windows with lower variance when timescales of correlations are shorter than the trial. Defined as the integral of the CCG over the integration time window, normalised by the geometric mean of the integral of the ACGs over the same period:

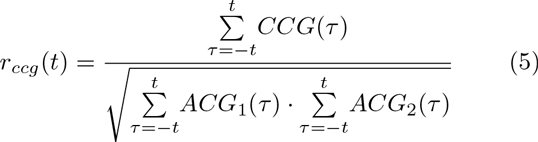

CCG and ACGs were corrected by the shu.e predictor. It can be shown that when considering the same time window *r_ccg_* is equivalent to spike count correlation (Bair et al., 2001).

Synchrony measures the tendency of neurons to fire with short and consistent delay between them. It was characterised using *A*_*ccg*_ (Smith and Kohn, 2008): the area under the jitter-corrected CCG (slow oscillations and stimulus locked variations are removed before integrating),with a 50 ms jitter window. The integration was performed between [-10,10] ms, in order to measure tightly coupled activity between pairs of neurons while removing loosely coordinated activity:

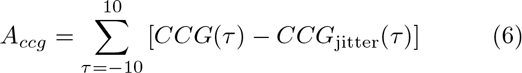
*A*_*ccg*_ allowed us to measure synchrony between pairs of neurons, which could be due to common synaptic inputs or synaptic connections. The correlation between synaptically connected pairs and synchrony was therefore studied.

Information carried by neurons about the stimulus was calculated using the software provided by Ince et al. (2009). We computed *I*_*sh*_ and used the NSB estimator (Nemenman et al., 2001) to compensate for sampling bias (the bias due to the limited number of trials available under experimental conditions). Information carried by single units was calculated using a variable time window to evaluate the effect of timing on the information conveyed by the evoked response. On each trial the neuron’s response corresponded to a vector with the number of spikes fired on each time bin. The binning ranged from 40 ms (full time window, equivalent to spike count analysis) to 5 ms (at which the information carried by single units started to saturate). Information analysis for triplets and pairs of neurons was done with a time window of 40 ms (spike count) and 10 ms (a time window that allowed a reduced number of possible response symbols, yet keeping temporal information). To reduce the number of possible responses spike counts were thresholded at the 95th percentile for 40 ms time windows and only allowed to be 0/1 for 10 ms (4) time windows. Response of pairs and triplets was simply the concatenated vector response of the individual neurons. Information gain for pairs and triplets was calculated on each case by subtracting the information obtained from the joint response minus the sum of information obtained by individual responses.

### 2.4. *Local field potential characterisation*

LFP conveys information about synaptic currents (Buzsaki et al., 2012) and its tuning can be then compared to tuning seen on the local neuronal population. We included the 4-40 Hz frequency range which has been shown to convey tuning information (Eggermont et al., 2011). LFP baseline was calculated over the time window minus 12 to 0 ms relative to stimulus onset. The baseline LFP was then integrated over the 0 to 40 ms time window to obtain the evoked LFP for each frequency. The best frequency for the LFP was defined as that with the largest integrated value. LFP latency was measured at this frequency from onset to peak response.

## 3. Results

### 3.1. *The local columnar population presents similar response properties, but large differences are possible*

We recorded from 231 neurons from 12 mice. Of these, 82 neurons demonstrated frequency tuning. Our findings (values reported as mean ± s.e.m unless otherwise stated) are in agreement with previous reports (Rothschild et al., 2010; Hromádka et al., 2008) of sparse responses in anaesthetised mouse auditory cortex. Recorded neurons were classified as narrow spiking (24.7 %) or broad spiking (75.3 %). Mean spontaneous firing rate was 1.1 ± 0.21 spikes/sec, while mean evoked firing rate was 2.7 ± 0.47 spikes/sec. 90% of the cells had evoked firing rate below 7.1 spikes/sec. These low firing rates are in agreement with previous reports (Hromádka et al., 2008; Bandyopadhyay et al., 2010; DeWeese et al., 2003). We found the mean preferred frequency across neurons to be 16.7 ± 1.4 kHz, mean latency to be 26.8 ± 0.94 ms, mean LFP latency 27.1 ± 1.1 ms and mean bandwidth 6.3 ± 0.64 kHz. These values are in accordance with previous reports showing a non-homogeneous representation of frequencies in primary auditory cortex, with a peak at around 22 kHz (Rothschild et al., 2010), and the mean delay in layer five measured to be around 21 ms by Sakata and Harris (2009).

**Figure 1:**
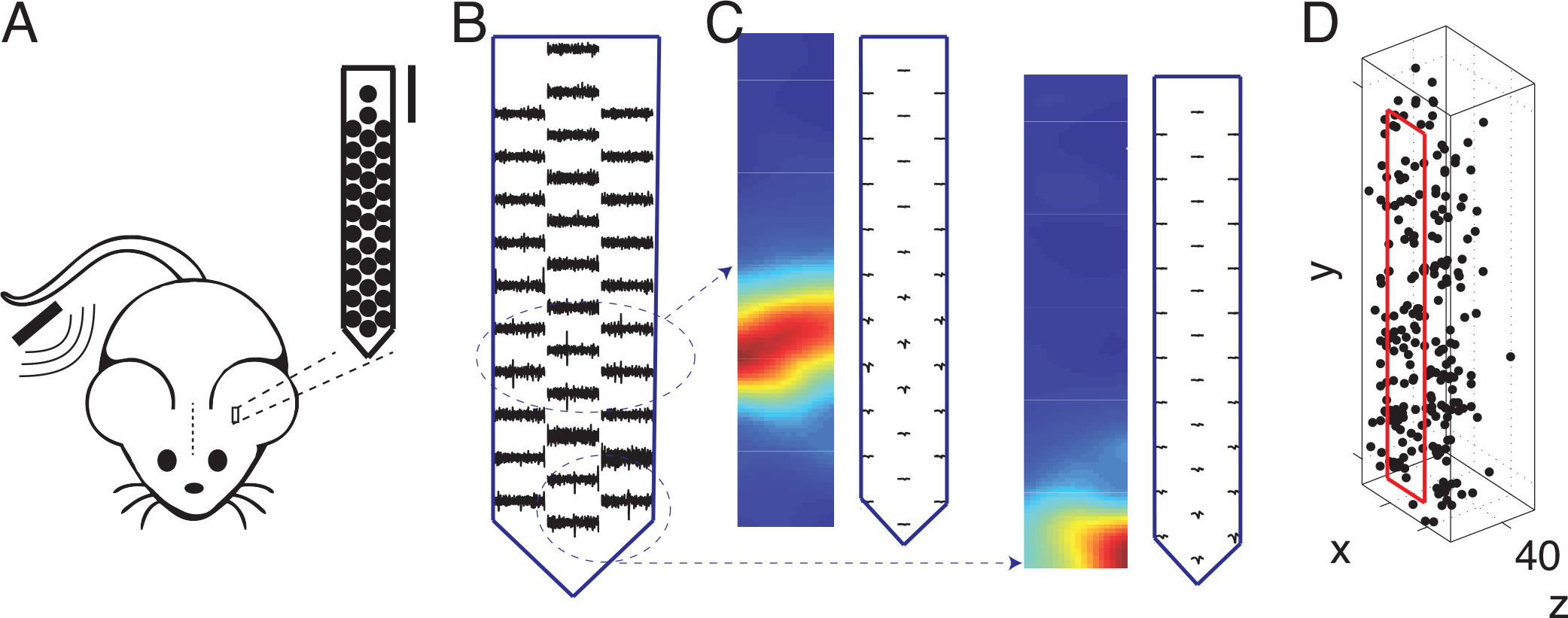
Extracellular recordings with high-density electrode array targeted microcolumns in A1. (A) We targeted layer five of contralateral primary auditory cortex, inserting the probe parallel to midline to maximise rostral to caudal coverage and perpendicular to the brain surface. (B) A typical recording, here highlighting two isolated neighbouring units. (C) Single units from (B) illustrated independently, showing the average spike waveform measured on each channel and the corresponding heatmap depicting strength of the signal across channels. D) Three dimensional plot showing estimated location of the soma of all single unit on the dataset

In Fig. 2 we show data from two representative population recordings. In the first column we see the estimated location of each neuron on the xy plane relative to the probe, and colour coded their preferred frequency. We can appreciate that in general they tend to prefer nearby frequencies. Qualitatively we can see from the tuning curves shown on the second column that having similar preferred frequency, their particular profile across frequencies can still differ greatly. From the third column, showing raster plots of their evoked responses across trials, we can see that their temporal response profile can also be very dissimilar and within each recording we found a variety of temporal response profiles. These differences on their temporal profile are an indication of the different processing done by each neuron and their temporal profile likely determines temporal windows relevant for downstream neurons.

**Figure 2:**
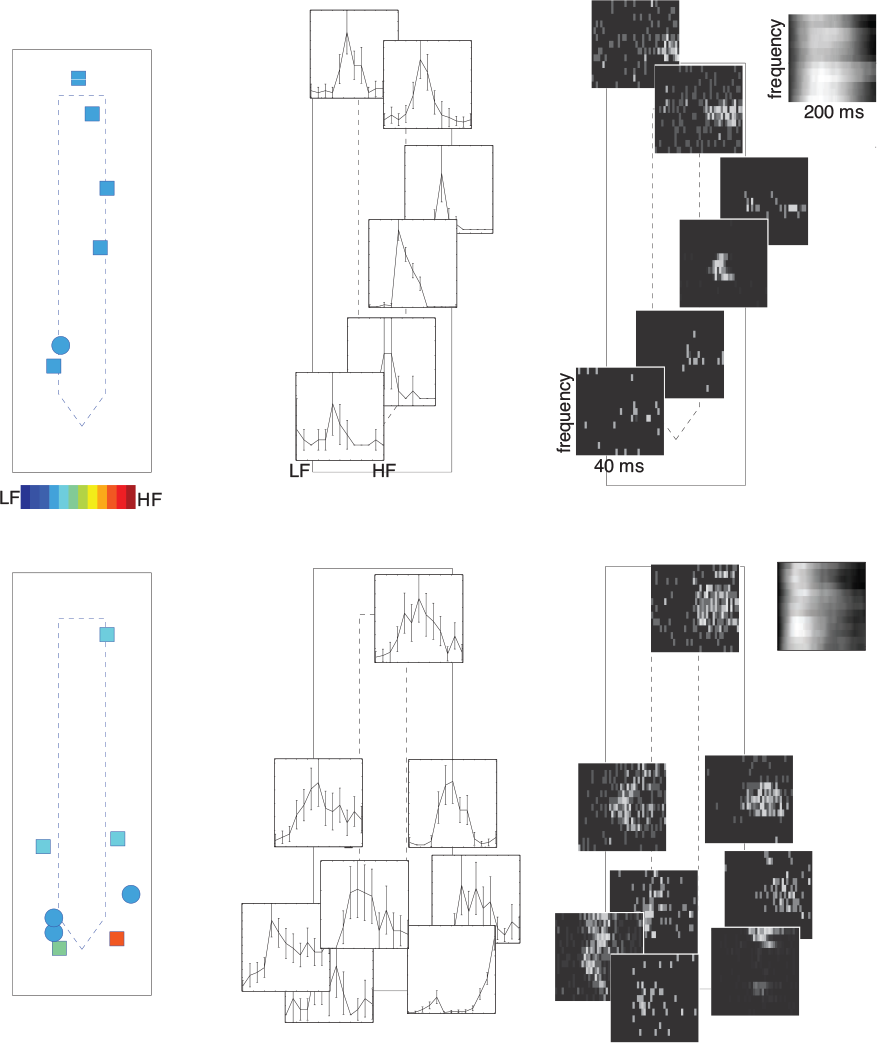
Within micro columns we found a variety of response profiles with a tendency to share similar preferred frequency. For two recordings we show details of the response to our stimulus **left** xy plane showing the edges of the probe and the location of each neuron relative to the probe, the colour corresponds to its preferred frequency **middle** for the same units we show the tuning curve (frequency versus probability of firing) right raster plots of the neurons’ response after stimulus onset (40 ms) across trials (differing on frequency). Top **right** corner shows the temporal profile (over 200 ms after stimulus onset) for the LFP across frequencies.

We found that in deep layers of primary auditory cortex, neurons tend to share similar response properties. When comparing simultaneously recorded neurons, the mean difference in properties exhibited by simultaneously recorded neurons was small compared to the full range spanned by the responses, as seen in table 1 (each column listing mean, s.e.m, 95th percentile and maximum difference respectively). A fractured representation was present in some recordings, as can be seen from the Voronoi diagrams in Fig. 3A where we can appreciate smoothness which is fractured by sudden changes. While the tendency was for neurons to present similar preferred frequency as the LFP (Fig. 3B) and for pairs of neurons to share similar response properties, some pairs displayed quite dissimilar properties (lying at the opposite ends of the spectrum), as can be seen in the histograms of Fig. 3C and last column of table 1 which lists the maximum difference measured between pairs of neurons.

**Figure 3:**
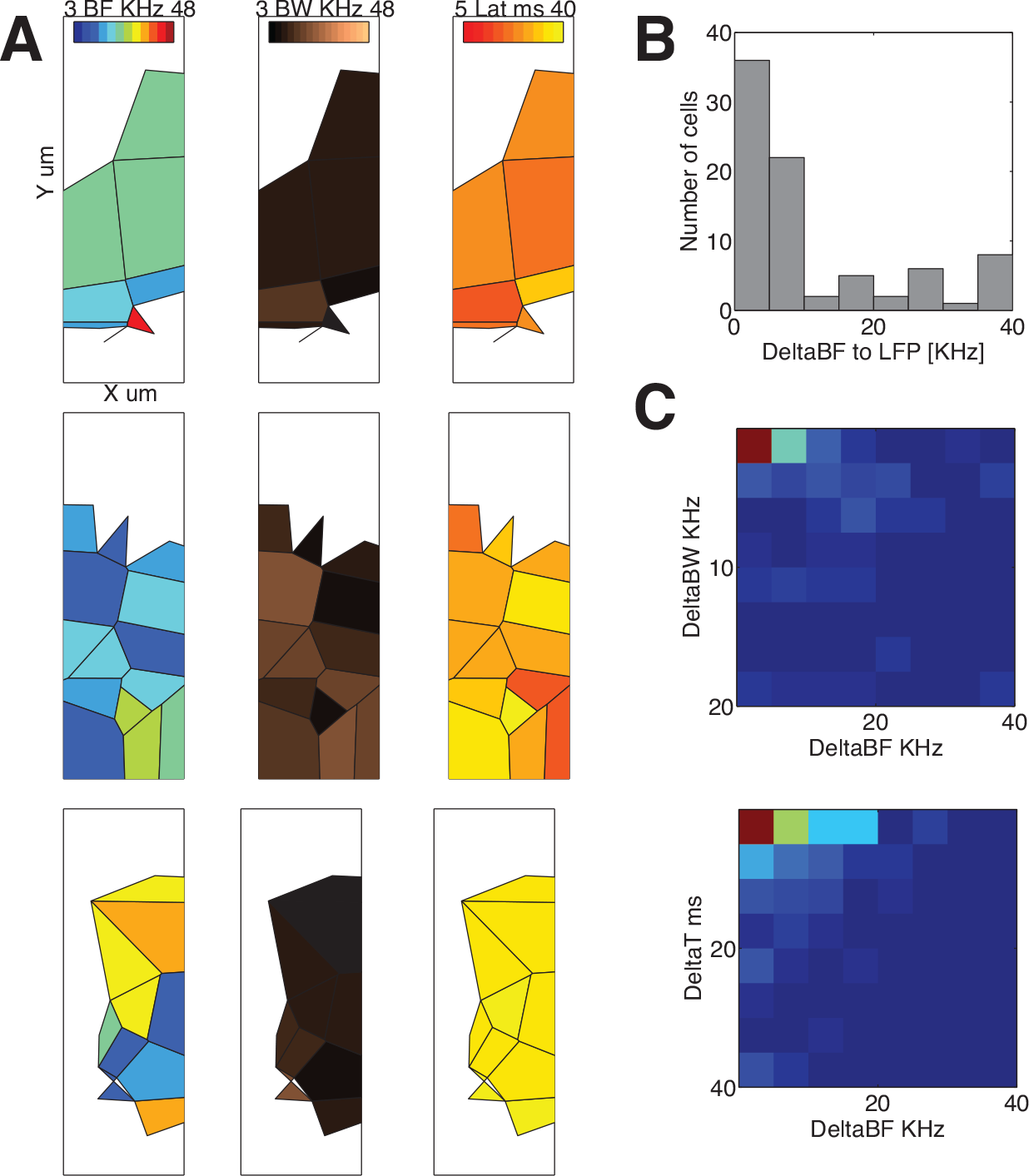
Microcolumns exhibited fractured representations of response features. (**A**) For three recordings we show Voronoi diagrams colour coded in terms of preferred frequency, bandwidth and latency from onset. Here we can see that the smoothness is broken by sudden differences between neighbouring neurons.(**B**) Neurons tend to have similar preferred frequency as the LFP, but large deviations from it were also possible. (**C**) Histograms of difference in latency and bandwidth versus difference in preferred frequency. Neurons tend to have similar properties, but large differences were also possible.

**Table 1:**
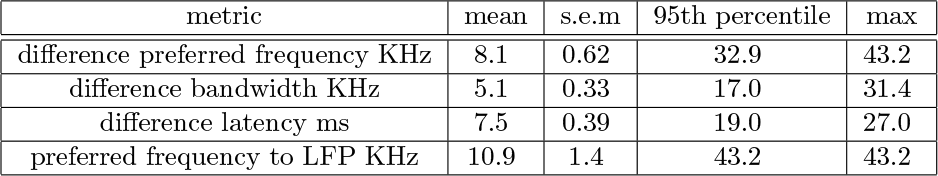
Differences in response properties between simultaneously recorded pairs of neurons.

| metric | mean | s.e.m | 95th percentile | max |
| --- | --- | --- | --- | --- |
| difference preferred frequency KHz | 8.1 | 0.62 | 32.9 | 43.2 |
| difference bandwidth KHz | 5.1 | 0.33 | 17.0 | 31.4 |
| difference latency ms | 7.5 | 0.39 | 19.0 | 27.0 |
| preferred frequency to LFP KHz | 10.9 | 1.4 | 43.2 | 43.2 |

Difference seen in preferred frequency, bandwidth, delay and best frequency to **LFP**.

**Table 2:** Changes in difference of response properties over distance.

| metric | under 100 μm | above 150 μm |
| --- | --- | --- |
| difference preferred frequency KHz | $8.6 \pm 0.87$ | $5.2 \pm 0.89$ |
| difference bandwidth KHz | $5.2 \pm 0.44$ | $5.4 \pm 0.78$ |
| difference latency ms | $7.0 \pm 0.48$ | $7.8 \pm 0.92$ |

We compare for all response properties (preferred frequency, bandwidth, latency) the mean±sem for neurons located close (under 100 *μm* appart) and far (over 150 *μm* away). Differences were not statistically significant

The similarity in terms of stimulus preference was quantified by the signal correlation, which was high on average (0.58 ±0.023), however this large signal correlation does not reflect differences on the temporal profiles of the cells seen on the raster plots in Fig. 2 which might indicate that a more complex stimuli would yield a lower signal correlation. A relatively large proportion of pairs of neurons exhibited quite similar response properties at all separations up to several hundred microns apart, as indicated by the large fraction of neurons at all separations with signal correlation approaching 1.0 in Fig. 4A. Nevertheless, there was also a significant tail of neurons showing quite dissimilar tuning (neurons with signal correlation close to zero in Fig. 4A). There was no significant linear correlation between signal correlation and distance (Pearson r = −0.024), with pairs spanning the full range of signal correlation at all distances.

There was no significant dependence of difference in response properties and distance between cells, as shown in Fig. 4B-D. When comparing the mean for neurons located closer than 100 *μ*m apart versus neurons more than 150 *μ*m apart from each other, as listed in table 2, we found that: for differences in preferred frequency, although apparently decreasing with distance, the change was not statistically significant(Mann-Whitney U-test); differences in bandwidth did not show change with distance; differences in latency, although apparently increasing on average with distance, this change was not statistically significant.

**Figure 4:**
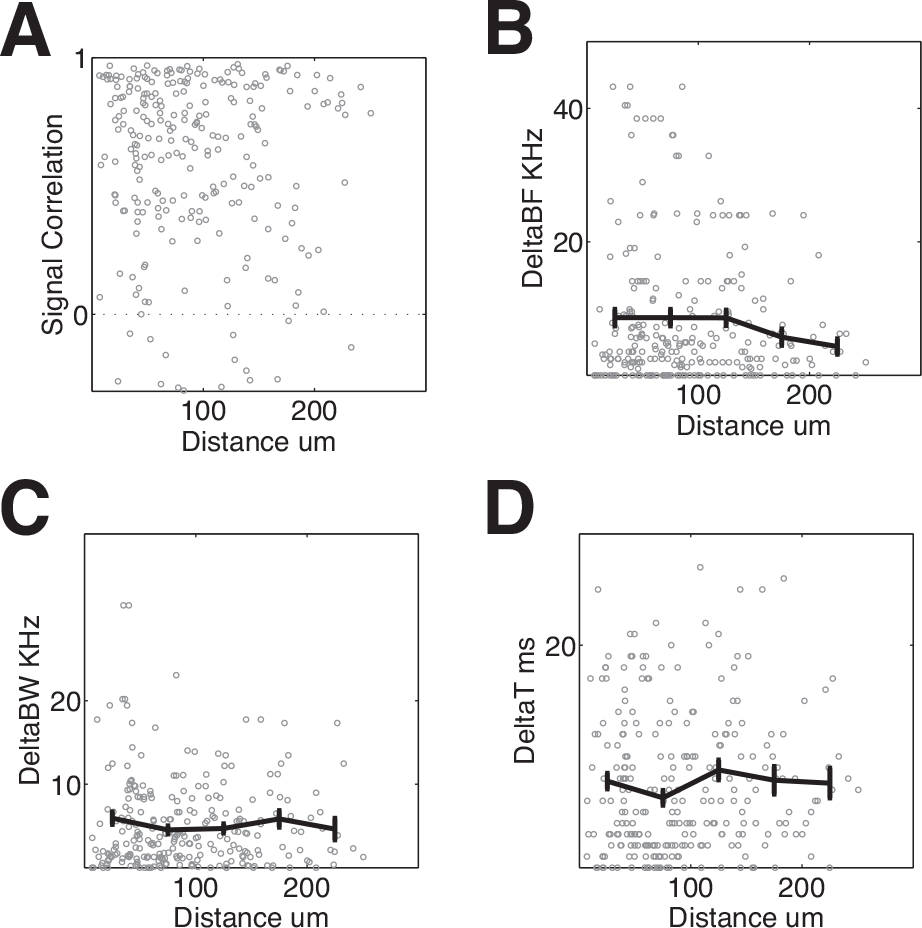
Differences in response properties versus distance showed no significant relationship. (**A**) Signal correlation between simultaneously recorded pairs showed no significant trend with distance (bars represent mean±sem, as in all other raster plots), at all distances signal correlation spanned the full range. (**B**) Difference in preferred frequency versus distance. Larger differences were more common at short separations, but the mean difference showed no significant correlation with distance. (**C**) Difference in bandwidth also showed no significant correlation with distance and the full range was spanned at all distances. (**D**) Difference in peak latency versus distance. There was no significant linear correlation and small and large differences were found at all distances.

### 3.2. *Correlation and synchrony decay with distance and during evoked activity*

Noise correlation measures the tendency of the trial to trial fluctuations of neuronal responses to the same stimulus to correlate with each other. This can be due to slow oscillations (such as the changes seen during up and down states) or faster synchronous activity generated by shared inputs unrelated to the stimulus. We measured the noise correlation between pairs of neurons over the 40 ms time window used above to compute the signal correlation. Noise was positively correlated with signal correlation as seen in Fig. 5A, meaning that neurons that share more stimuli related inputs also share more substrate inputs. Noise correlation was not heavily dependent on distance as seen in Fig. 5B, but larger values only occurred at shorter distances. The 95th percentile could be bound by an exponential of decay constant 169.7 *μm,* which implies that over this distance the largest values of noise correlation are reduced on approximately 60%.

Our population average noise correlation of 0.057 ± 0.0051 is in agreement with Sakata and Harris (2009), but is 4 times smaller than that reported by Rothschild et al. (2010). This could be attributed to the time window used to compute noise correlation, since as seen in Fig. 5C our correlation saturates at similar mean values seen in (Rothschild et al., 2010) when increasing the time window to match the 200 ms used in that study. However, large windows are unlikely to be relevant when considering processing of information done by downstream neurons (Cohen and Kohn, 2011).

The noise correlation *r*_*ccg*_ observed during spontaneous activity was greater than that observed during evoked activity (as seen in Fig. 5D for varying integration windows from 0 to 200 ms). Mean *r*_*ccg*_ of 0.13 ± 0.0063 for spontaneous, compared to 0.097 ± 0.0050 for evoked at T = 40 ms which corresponds to the time window used to characterize tuning and signal correlation; p=0.00014 at 5% significance, n=263.0 pairs of tunned neurons, Wilcoxon rank sum test). We interpret this as de-correlation of the local population during sensory stimulation.

To study changes in synchrony (correlation at very short timescales), we measured the area under the crosscorrel-ogram *A*_*ccg*_. We observed a greater synchrony in spontaneous as compared to evoked activity: 0.029 ± 0.00048 and 0.024 ± 0.00033 coincidences per spike respectively. This is reminiscent of similar results observed in primate visual cortex (Kohn and Smith, 2005). Synchrony during evoked activity was 42.6 % (best fitting line) of the levels of synchrony measured during spontaneous, this compares to 70.1 % seen in correlation, the comparison of spontaneous versus evoked can be seen qualitatively in Fig. 5D and E. This means that the loss of synchrony was more significant that changes at slower timescales. Therefore we can conclude that different network configurations are active during evoked and spontaneous activity, driving neurons on the local population to different correlation and synchrony regimes, this particular stimulus did not change significantly slower oscillations but had a significant impact on synchrony. In terms of its dependance on distance, the largest recorded values of synchrony decayed faster during evoked activity than during spontaneous, as can be seen from Fig. 5F where a exponential was fitted to the 95th percentile with decay constants of 317.1 and 446.8 *μm* for evoked and spontaneous activity respectively, which could imply overlaping network configurations that differ on their physical coverage being more confined during evoked activity.

**Figure 5:**
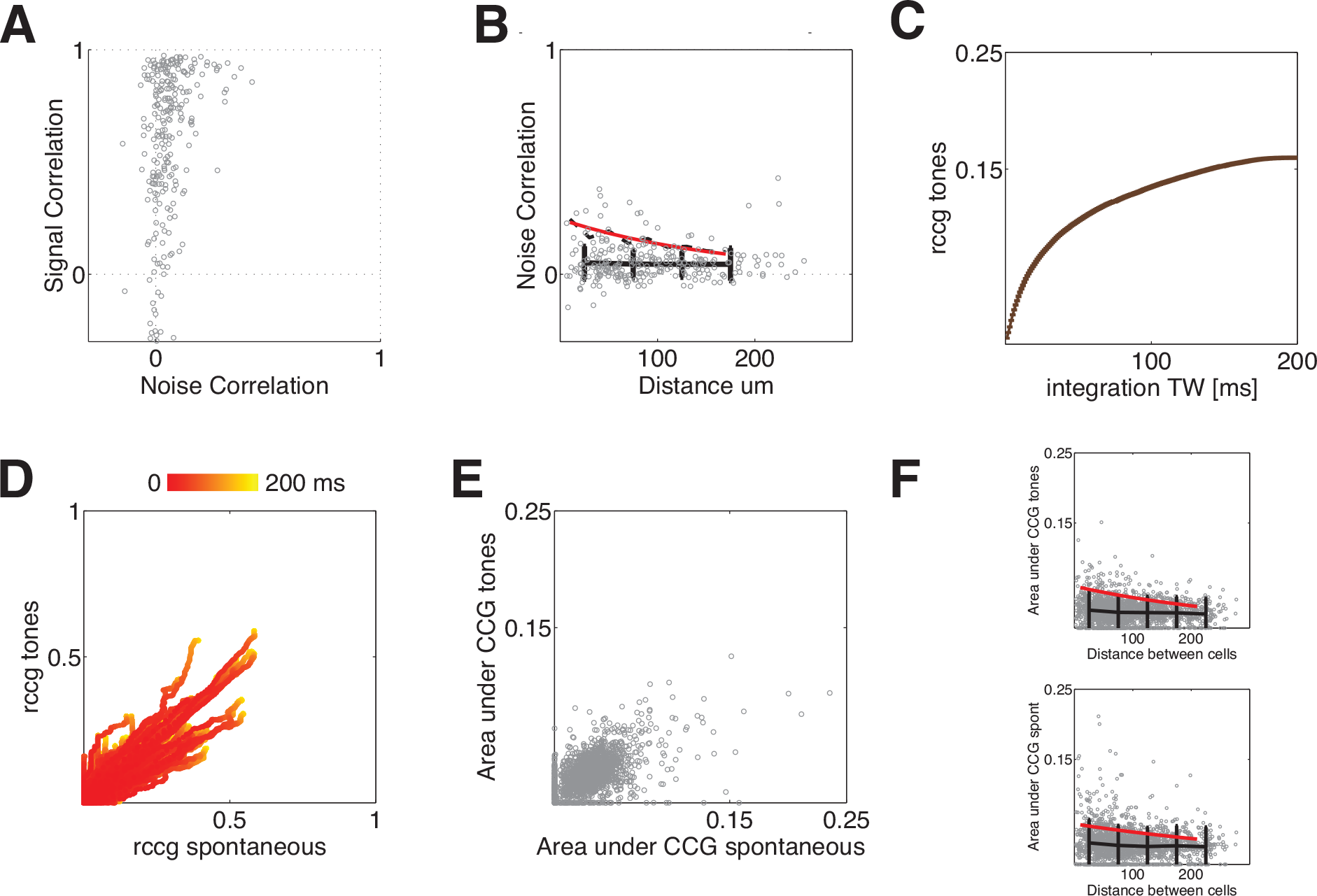
Low average noise correlation and stimulus induced decorrelation. (**A**) Signal and noise correlation were positively correlated.(**B**) Noise correlation was lower on average than previously shown and showed no significant dependence with distance, but larger values were more common at shorter distances. Black bars correspond to mean±sem, segmented black line 95th percentile and red line exponential decay fitted to it.(**C**) When analysing the timescales of correlation (*r*_*ccg*_) we found that saturation values are similar to values previously reported, although such large time windows are unlikely to be relevant for upstream neurons. (**D**) *r*_*ccg*_ measured during evoked versus spontaneous activity, colour coded by time window. (**E**) Synchrony (A_ccg_) measured during evoked versus spontaneous activity. (**F**) Synchrony versus distance between neurons, measured during evoked (top) and spontaneous (bottom). Black bars represent mean±sem, red line the exponential decay fitted to the 95th percentile

### 3.3. *Groups of neurons carry redundant information on average, but high information gains are achieved on similarly tuned pairs*

We calculated the information carried by single units, pairs and triplets of neurons. Single unit analysis was done over varying size of binning time window, starting from 40 ms (equivalent to evaluating the information carried by spike count) we decreased the time window to see the impact of timing on the information carried by each neuron. From Fig. 6A we see that information started to saturate at 5 ms when the average information carried per neuron was 4.5 ±0.48 bits/sec and the average information carried by a single spike was of 0.96 ±0.073 bits (Fig. 6B). The distribution of information carried by single spikes was not homogenous as seen in Fig. 6C, the distribution was skewed and only a tail of neurons carried a large amount of information per spike. There was no difference between broad spiking and narrow spiking neurons (mean: 0.95 ±0.079 bits/spike versus 0.98 ±0.15 bits/spike, p=0.80, Mann-Whitney U-test)

We then measured the information gain obtained when considering the response of ensembles formed by pairs and triplets of neurons. The gain corresponds to the difference between the information obtained from the ensemble minus the sum of informations obtained when considering each neuron individually. This is identical to the percentage synergy measure used in some previous studies e.g. (Panzeri et al. (1999); Montani et al. (2007)). We studied information using a 40 ms time window (information carried by spike count) and 10 ms time window (a shorter time window produces a response space too large in order to estimate probabilities), by comparing both cases we can study how sensitive to timing is the information carried by the ensembles.

In Fig. 6D, we show the information carried by pairs versus information carried independently by the constituent neurons. Comparing the 40 ms binning (left) with the 10 ms binning (middle) we can appreciate that more pairs of neurons deviate from a full independent regime (diagonal) when timing is included on the vector response. This can be also seen from the information gain histograms (right panel) where we see that the spread of the fitted gaussian (red line) is larger for the 10 ms window (continuous line) and there is also an increase in the number of pairs with high information gain. While the maximum absolute information gain with a 40 ms window was 4.4 bps, the maximum gain with a 10 ms window was of 4.9 bps. In the mean time, the maximum relative gain for each time window was of 62.6 % and 231.3 % respectively. 3.8 % of the pairs had an information gain over 50% when using a 10 ms window, these pairs had on average a signal correlation of 0.76 ±0.066, higher than the average for all pairs (Mann-Whitney U-test, pval=0.048, 5% significance). These results tell us that while spike count does not have a large impact on information gain, when considering timing on the responses there is a gain for ensembles of neurons which have a high signal correlation corroborating the richness of the responses across the population despite their tuning similarity. On average however pairs of neurons carried redundant information as seen from the negative mean on the information gain histograms.

From Fig. 6E we see that a similar picture can be extracted for triplets of neurons. The relative information gain when using a 40 ms window was smaller than when using a 10 ms window (10.6 versus 5.5 bps of absolute gain and 43.3 versus 138.3 % of relative gain). Therefore, maximum relative gain for triplets was smaller than for pairs and the percentage of triplets with information gain above 50% was much lower than for pairs: 0.54 %. Also the percentage of triplets that showed synergy (gain of information) was smaller than the percentage of synergistic pairs (24.8 versus 39.8 %).

**Figure 6:**
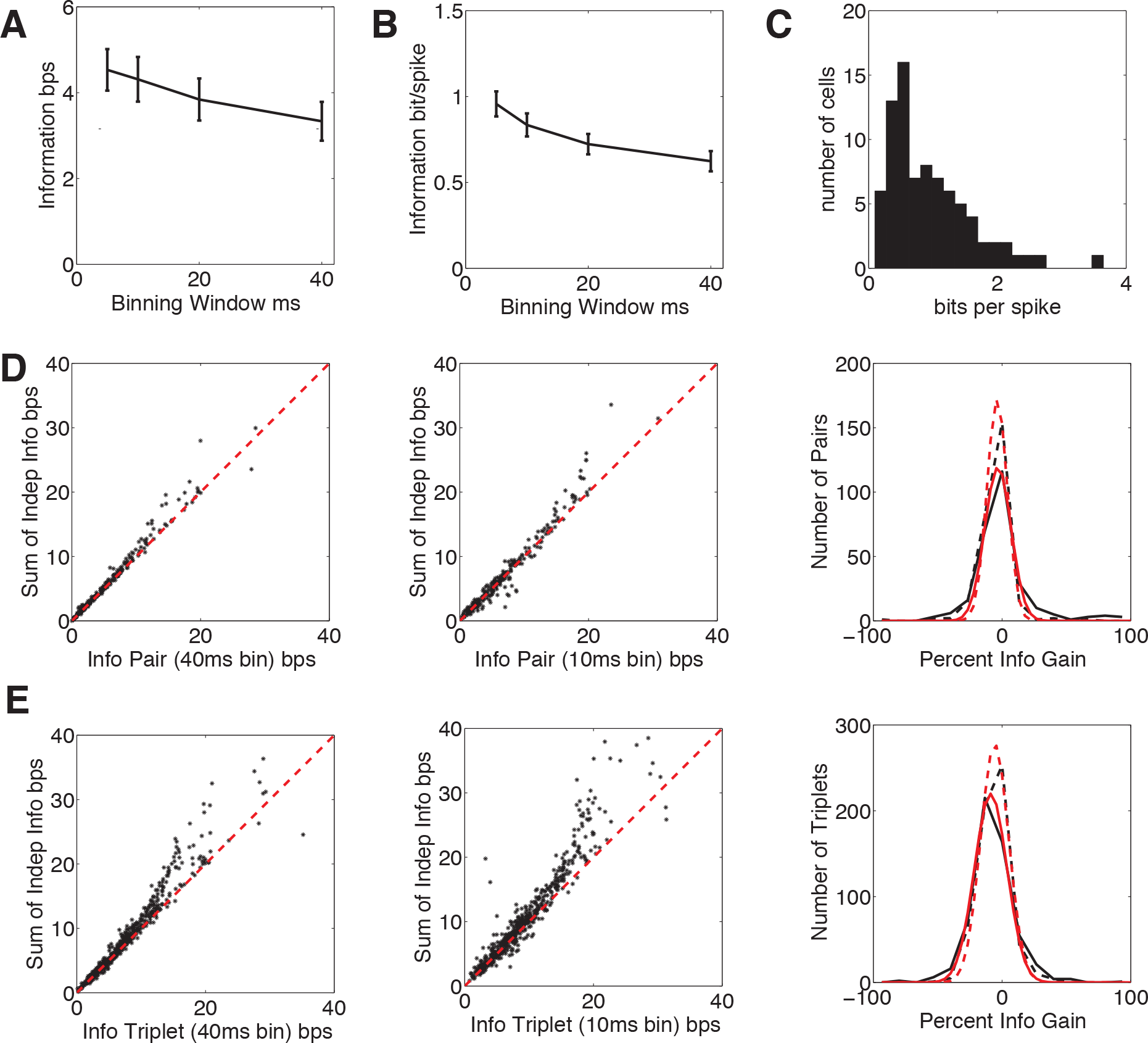
Information carried by single units, pairs and triplets. (**A**) Information in bits/sec (black bars represent mean±sem) carried by single units as a function of the binning window used to construct the vector response. (**B**) Information in bits/spike (mean±sem) also as a function of the window size. (**C**) Histogram of information in bits/spike carried by recorded neurons. (**D**) Rasters showing the information carried independently by neurons on a pair and the information carried by their joint response vector. Values deviate more from a fully independent regime (diagonal, red line) when including timing effects (10 ms binning window, middle) than when using rate code (40 ms window, left). This can also be seen on the histograms of percent information gain (percentage synergy) in the last column where black lines correspond to recorded distribution, and red line to a gaussian fit, larger gains are only present for the 10 ms binning window (continuous lines) compared to 40 ms window (segmented line). On average the population operated in a redundant regime (negative information gain, mean value for the normal distribution). (**E**) Results arranged as in (D), but for triplets. Similars conclusions can be drawn from them. Larger gains were achieved when including timming information (10 ms window, right) and also a larger spread on the distribution of information gains. Again for triplets the population on average operated on a redundant regime.

### 3.4. *Local connectivity is highly non-random*

The role that neurons play within microcircuits is determined both by their intrinsic properties and by feeding connections or interactions with other neurons. Previous studies have highlighted the importance of local connectivity, since it is expected that responses to stimuli are shaped by sensory related input (feedforward from thalamus in the input layers and upper layers in the case of layer V) and local interactions which in general have been assumed to further shape functional response. We therefore characterised connectivity for local populations of layer V neurons, in order to elucidate the role that these connections play.

Putative monosynaptic connections were detected based on crosscorrelograms. From our population of 231 neurons we found 21 to be exciting other neurons, with 20 of them corresponding to broad spiking neurons (putative excitatory neurons) and one possibly being excitatory with narrow spike waveform. The excited neurons (18) included both broad and narrow spiking (putative inhibitory) neurons: 10 and 8 neurons respectively. Considering that only nearly 20% of recorded cells are inhibitory (Meinecke and Peters, 1987; Xu et al.,2010), our results are consistent with a previous report that showed that probability of connection was higher on inhibitory neurons compared to excitatory, Yoshimura and Callaway (2005). This classification can be seen graphically in Fig. 7A left, were we can appreciate the correspondence between the excitatory and broad spiking class. In Fig. 7A right panel we can see that most of the neurons exciting others had a short refractory period and high burst index, which might suggest that they are large pyramidal cells from layer V, whose firing pattern matches that previously described by Christophe et al. (2005). Large pyramidal cells of layer V are known to be highly interconnected (Thomson and Lamy, 2007), in agreement with our observations.

In our dataset we found a low probability of connection: 0.0074, which matches low connection probabilities reported elsewhere (Song et al., 2005; Bartho et al., 2004). Despite this low probability, motifs of highly interconnected neurons were common, triplets of connected neurons occurred in 7 of the 10 recordings where synaptic connections were detected despite their low probability of occurrence: 0.00005. This result agrees with a previous in-vitro connectivity characterisation of layer V pyramidal neurons (Song et al., 2005), and is suggestive of highly nonrandom connectivity, with neurons forming sub-networks instead of loose pairs of synaptically connected neurons. The mean signal correlation between synaptically connected pairs was higher than the average signal correlation, but the difference was not statistically significant (Mann-Whitney U-test): 0.73 ± 0.081 versus 0.58 ± 0.023, respectively. The mean noise correlation was higher (pval=0.024, Mann-Whitney U-test) on average between connected pairs than the mean among all pairs: 0.14 ± 0.041 versus 0.057 ± 0.0051 see (Fig. 7D). A more complex stimulus might enhance differences on signal correlation while mantaining levels of noise correlation.

Synaptically connected pairs of neurons were on average synergistic when using a 10 ms binning window for the response vector (i.e. including timing effects), having on average 18.9 ±12.4 percent of information gain. The maximum gain among these pairs was of 86.0 % and only 2 pairs carried redundant information (negative information gain). In Fig. 7C we show an example of a triplet formed by a pair carrying redundant information (−18.2 % information gain, bottom row) and the other being slightly synergistic (3.5 % gain, top row), seeing the response pattern on these neurons across trials it would be interesting to explore the functional role these kind of synaptic connections play.

**Figure 7:**
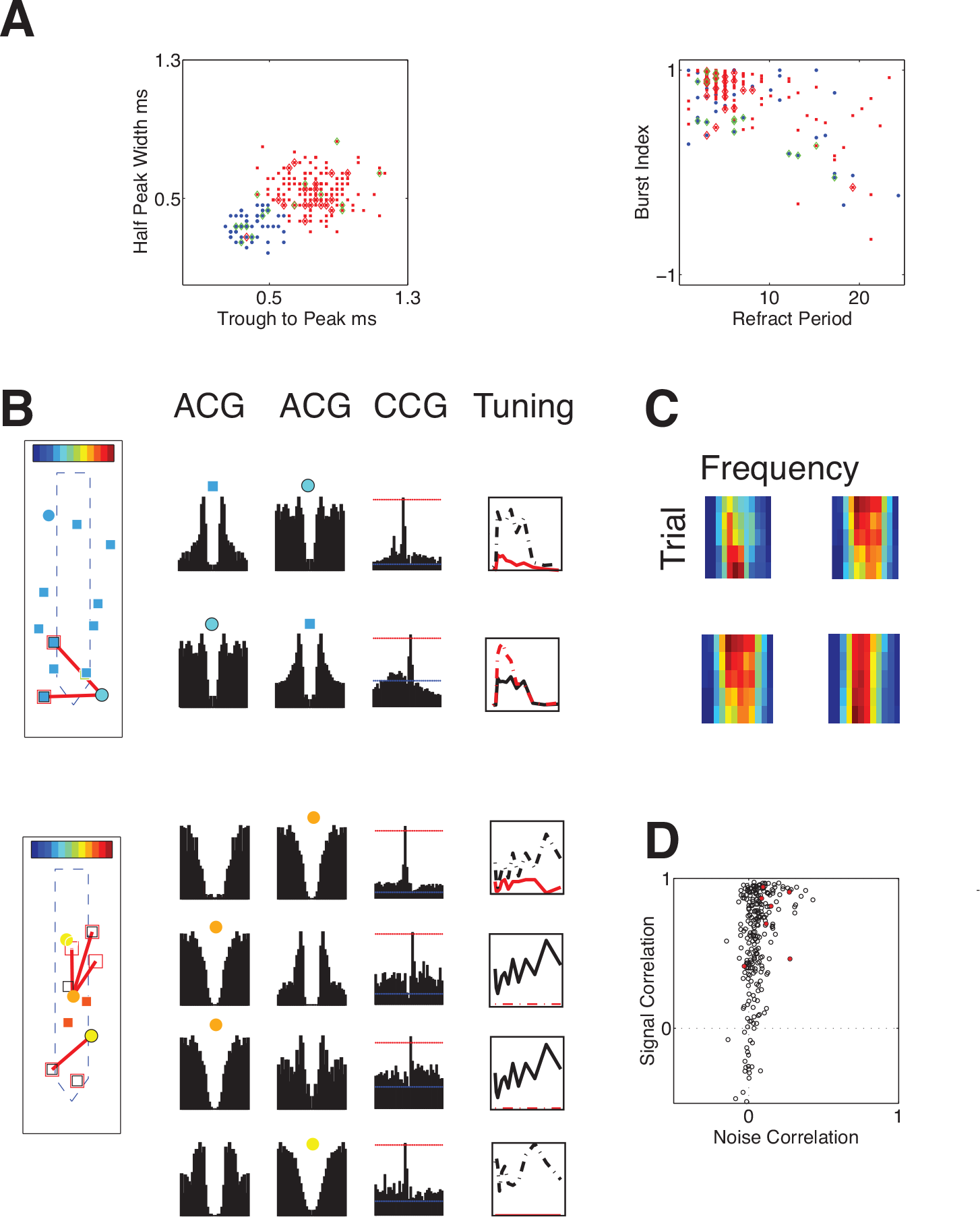
Highly non random connectivity. (**A**) **left** Neurons were classified as broad (red square) or narrow spiking (blue circle). Detected monosynaptic connections are also indicated here, excitatory neurons are indicated by a red diamond and excited neurons by a green diamond. **right** Characterisation of the firing pattern allowed to confirm classification of excitatory neurons as bursty neurons (putative large pyramidal neurons from layer 5).(**B**) Example of synaptic connections detected, for each the autocorrelogram of each neuron and crosscorrelogram are depicted along with tuning curves. Neurons are colour coded according to preferred frequency. (**C**) Example of synaptic pair corresponding to pair on adjacent **B** plot. The pair on the top row was slightly synergistic while the bottom was redundant (only 2 synaptic pairs carried redundant information). (**D**) Signal versus noise correlation for all (black circle) and synaptically connected (red asterisk), only difference in mean noise correlation was significantly different between both groups.

The mean separation along the rostro-caudal (X) axis between synaptically connected neurons was 28.3 μm, while the mean vertical (Y) separation was 48.9 *μ*m. The mean total distance was of 61.0 *μ*m. 90 % of detected pairs were located at less than 44.1,99.1 and 105.8 μm along the X, Y axis and total distance. These measurements are in agreement with Song et al. (2005), where the authors report for pyramidal neurons of layer V that 82% of the connections are located less than 50 *μ*m apart, with the remaining 18% being separated by less than 110 *μ*m. Our findings also agree with a connectivity study (Levy and Reyes, 2012), of layer 4 of auditory cortex reporting that the probability of connection is well fitted by a gaussian with a spread of 85-114 *μ*m, implying that the probability of synaptic connections decays as we move away in the coronal plane. Minicolumns have been proposed as a functional unit, with several mini columns interacting and being synaptically connected (Krieger et al., 2007), on that study synaptic connections were equally probable between neurons on the same bundle versus neurons belonging to neighbouring bundles. In contrast in our dataset we found that most of the connections, 87.5 %, were between neurons located over 10 *μ*m appart along the horizontal axis (rostro caudal) with 41.7 % of connections being between pairs located between 30 and 40 *μ*m. Although the columnar distribution in our dataset was not determined, given the columnar distribution of the electrodes themselves, we would argue that our findings are in agreement with the multi minicolumn interaction theory.

We measured how synchrony changed on spontaneous versus evoked activity. In the previous section we showed that for the whole population synchrony during evoked activity was a 42.6 % of that seen during spontaneous. For synaptically connected pairs we found this to be 62.6 %. This means that even for monosynaptically connected pairs, synchrony decreases when going from spontaneous to evoked activity, although the decrement is smaller than for the general population of neurons. Changes in synaptic strength have been reported previously (Fujisawa et al., 2008) and could imply different subnetworks operating on a local columnar population.

## 4. Discussion

Recent studies (Rothschild et al., 2010; Bandyopadhyay et al., 2010; Hackett et al., 2011; Guo et al., 2012) have examined the response properties of the superficial layers of A1. They have reported tonotopy when recording multi unit activity and LFP, but fractured tonotopy at the single neuron scale, with neighbouring neurons exhibiting very different frequency preferences. Tonotopy has been reported to be stronger in the thalamo-cortical recipient layer, and degraded in downstream layers. In this study we examined tonotopic structure in local neuronal populations in the main output layer of A1 (layer 5) with the aim of characterizing their response properties.

Our study, the first use of dense columnar recording technology to map tonotopy in mouse auditory cortex, revealed fractured tonotopy at the level of single neurons: neighbouring neurons tend to share frequency preferences, but with occasional large differences in tuning. Previous multi-electrode array electrophysiological studies have shown tonotopy at a scale of hundreds of microns (e.g. (Guo et al., 2012) which used a sparse array and (Hackett et al., 2011) which used a high density linear array shifted in position over hundreds of *μ*m), but due to relatively sparse spatial sampling, these studies were unable to provide evidence of the detailed topographic organisation of auditory cortex on small volumes (From our estimated locations, the maximum scanned volume was of 0.0015 mm^3^, spanning 84.2 *μ*m rostro-caudal, 277.2 *μ*m vertical and 64.2 *μ*m lateral). Several recent studies, using two photon calcium imaging, found a high degree of heterogeneity in the tuning of spiking-related calcium signals at the single neuron scale (Rothschild et al., 2010; Bandyopadhyay et al., 2010). One possible reason for the difference between our finding and these studies may be that they are effectively sampling from a single coronal plane covering layers II/III, whereas we densely sampled in the columnar axis targeting only layer V. The groups of similarly tuned neurons we observed could have a physical correlate, for example micro columns or bundles (Maruoka et al., 2011; Krieger et al., 2007; Ohki et al., 2005), while being compatible with neighbouring groups exhibiting large differences in their preferred frequency as found by our study and the aforementioned imaging studies. The incidence of large differences in preferred frequency might be related to the angle of insertion, with slightly tilted angles increasing the probability of finding large differences between neighbouring neurons.

The specific pattern across frequencies varied substantially between neighbouring neurons, despite their tendency to operate on the same frequency band, we found a large variability within the local population. On average simultaneously recorded neurons exhibited similar preferred frequency and bandwidth, but large differences were possible at all separations between cells. This is also seen from the large variability in signal correlation found in the local population at all distances, with a higher mean signal correlation than found in the previous studies which made use of two photon microscopy (Rothschild et al., 2010; Bandyopadhyay et al., 2010). This difference in mean signal correlation could be explained by the columnar sampling of the population targeted by us as opposed to the spanning of larger surfaces in the coronal plane: in the former case, a larger amount of common input might be expected. The use of iso-intensity tones also increases the signal correlation since differences in intensity tuning are not considered. Despite the simplicity of the stimuli used, we can conclude that local populations in layer V constitute a set of complex and diverse filters. Further studies with complex stimuli might help elucidate how different spectrotemporal filters coexisting in the local columnar population complement each other.

The temporal response patterns seen on local populations were also diverse, despite their tendency to have similar peak latency from onset, with nearby neurons exhibiting different temporal profiles (for example on/off response, sustained, transient). This again tells us about the richness of response properties among the local population, with neighbouring neurons potentially performing very different processing tasks. The temporal profile determines the time windows and temporal dynamics with which neurons convey information about the stimuli, and is defined by both the neuron’s intrinsic properties, and those of upstream feeding neurons.

Noise correlation was congruent with previous reported values when analysing over similar time window. Noise correlation was found to decrease during evoked activity, therefore the stimulus used effectively decorrelated the activity of the local population. Synchrony was also found to decrease during evoked activity, which indicates that even neurons having a tighter connectivity see a change on their connectivity strenght when stimulated. Average levels of noise correlation and synchrony didn’ show a dependance with distance. However, the largest recorded values decayed with distance in both cases, noise correlation decayed faster than synchrony which in turn decayed faster during evoked activity compared to spontaneous. This indicates that networks directly involved on the transmision of evoked activity are more localized than networks sustaining spontaneous activity. The superposition of common inputs driving the local population (providing sub threshold oscillations, for example) and non random local connectivity with the presence of hubs of connected neurons, as seen in our recordings, would explain the faster decay with distance of noise correlation compared to synchrony. In support of this Yoshimura and Callaway (2005) showed that synaptically connected neurons do not always share common excitatory inputs, a fact which can be explained by complex fine-scale networks at a sub-columnar scale (Yoshimura et al., 2005; Yoshimura and Callaway, 2005). There could be overlapping subnetworks which might be activated under different stimuli conditions (Fujisawa et al., 2008) as hinted by the reduction in synchrony seen between synaptically connected neurons when switching from spontaneous to evoked.

When analysing the information carried by single neurons we found that the rate of transmision and the information carried by each spike grew as we decreased the binning window, which means that the code used by neurons in mouse A1 to transmit information has a resolution below 5 ms, or conversely, downstream neurons could potentially decode responses at at least this time scale. A more complex stimulus might yield an even higher gain as the window is reduced, as shown in (Liu and Schreiner, 2007) for information carried by primary auditory cortex neurons of mothers when stimulated with pup’s crying sounds. The distribution of information carried by single spikes was not uniform but skewed in agreement with previous observations from macaque auditory cortex (Ince et al., 2013).

Pairs of simultaneously recorded neurons were found to carry redundant information on average, but large gains (synergy) were observed for pairs of neurons when temporal information was included on the response vector, these levels of synergy were not seen when considering only spike counts. We found that the mean signal correlation among pairs which had a large synergy (above 50% information gain) was higher than the average signal correlation, which means that even when pairs showed similar response to the stimulus they still carried complementary information on the temporal profile of their responses. Also pairs of monosynaptically connected neurons were on average synergistic, which implies that despite their common activity due to direct connectivity they can still carry complementary information. However we found few examples of synaptic pairs operating in redundant mode and looking at their response it would be worth investigating their role, specially given their location on the main output layer of cortex.

Triplets of neurons showed a similar result, by having large synergy when temporal information was included on the response vector. However, the largest gain seen on triplets was lower than that measured for pairs and the percentage of triplets with large gains was smaller than the percentage of pairs. The percentage of triplets operating on synergistic mode was smaller than the percentage of synergistic pairs, this results concords with findings from rat motor cortex (Narayanan et al., 2005) which found around 20% of pairs to be synergistic, while 99% of larger groups (8 neurons) were redundant. We could expect to find more synergy and less redundancy when using a more complex stimulus, since a simple stimulus can increase the level of redundancy by unexploiting the encoding capacity of the neurons. Low levels of redundancy are also expected according to a study from cat auditory cortex (Chechik et al., 2006) which reported that redundancy decreases as we move along the auditory pathway. The analysis performed for groups of neurons rely on the assumptions done about the decoding scheme used by downstream neurons, such as not pooling spikes coming from different neurons, which might not be the case if the feeding neurons synapse on equivalent locations (see (Schultz et al., 2009) for a discussion of this issue in a quite different circuit, that of the cerebellum). Synergy can be increased by pooling if there is signal anticorrelation (Panzeri et al.,1999); conversely, for positive signal correlation, synergy is improved by considering each neuron independently. It is likely that both cases are produced by the connectivity patterns seen in cortex. There are also cases where redundancy is desirable, gaining reliability to the detriment of information rate (Barlow, 2001).

Modelling studies (Kaschube et al., 2010) have suggested that long range suppressive connections are in charge of shaping maps and the existence of pinwheels, while local connectivity might dictate arrangement in areas lacking orderly maps as is the case of mouse auditory (and visual) cortex. However, connectivity patterns and the role they play might differ across layers. A previous report from the upper layers of auditory cortex (Rothschild et al., 2010) explained their findings by a tonotopically arranged input (local neurons sharing common input) and random local connectivity. This would explain the decrease in correlation they observed as the distance in the coronal plane increases and the variability seen at short distances, which could be explained by overlapping sub-networks. The existence of such networks has been reported from studies in rat visual cortex (Yoshimura et al., 2005; Yoshimura and Callaway, 2005). In this study we found evidence for functional sub-networks in deep layer of A1, however the connectivity seen differed from random.

In layer V, we found that connectivity was highly nonrandom, and triplets of synaptically connected neurons were the most common motif, despite their extremely low probability of occurance, in agreement with a previous in-vitro study (Song et al., 2005). Columnar processing has long been assumed to be a hallmark of cortex, and a in-vitro study (Krieger et al., 2007) has suggested that an aggregation of mini columns might act as a single functional unit. Supporting this concept, we found that most monosynaptic connections were between neurons located on different vertical bundles (given by their separation along the rostro-caudal axis), the fact that synaptic pairs were on average synergistic supports the idea of functional aggregation of adjacent columns. Further studies with recording arrays spanning several mini columns may allow a thorough characterization of connectivity between them, and the determination of their functional correlates in-vivo.

## 5. Acknowledgements

We acknowledge the support of the Arcadia Fund (studentship to IDR), the Royal Society (Industry Fellowship to SRS), BBSRC grant BB/K001817/1 to SRS, and EU FP7 grant 289146 to SRS.

